# Two new gastropod holobionts from Atlantic hydrothermal vents showing different levels of endosymbiosis dependence

**DOI:** 10.64898/2026.10.05.756691

**Authors:** Pierre Methou, Xueying Nie, Ouafae Rouxel, Loic N. Michel, Magali Zbinden, Valérie Cueff-Gauchard, Johanne Aubé, Joan M. Alfaro-Lucas, André Ampuero, Marjolaine Matabos, Julia D. Sigwart, Cecile Cathalot, Thomas Giunta, Chong Chen, Marie-Anne Cambon, Jin Sun

## Abstract

Chemosynthetic symbioses underpin animal life at deep-sea hydrothermal vents, encompassing substantial variabilities in symbiont identity, metabolic capacity, and host dependence. Here, we characterise symbioses in two peltospirid snails from Mid-Atlantic Ridge vents, *Peltospira gargantua* and *P. smaragdina*, using 3D anatomical reconstruction, stable isotope analyses, fluorescence *in situ* hybridization, transmission electron microscopy, and metagenomics. Both species hosted abundant intracellular *Campylobacterota* within bacteriocytes of a hypertrophied oesophageal gland. Their dominant symbionts belonged to genus *Sulfurimonas*, but the two hosts were associated with distinct symbiont lineages. Symbiont genomes encoded carbon fixation through the rTCA cycle, sulfide oxidation, and Group I Ni-Fe hydrogenases, indicating combined thiotrophic and hydrogenotrophic capacities. Hydrogenase copy number varied across vent fields from one to three copies. The *P. gargantua* symbiont from the hydrogen-enriched Hydra vent field carried two divergent copies, including one with a truncated catalytic region, potentially reflecting local adaptation to vent geochemistry. Anatomical and isotopic comparisons further revealed contrasting host dependence, with *P. gargantua* having a proportionally larger oesophageal gland and ventricle coupled with a reduced radula, as well as a greater nitrogen-isotope enrichment between symbiotic and non-symbiotic tissues. The smaller oesophageal gland and larger feeding apparatus of *P. smaragdina* point to it being mixotrophic. Previously, all known peltospirid symbioses have partnered with *Gammaproteobacteria*, and we expand the known diversity of peltospirid symbioses. Our findings underscore how closely related hosts can exhibit different degrees of dependence on symbiosis, and that mixotrophic symbiosis may be prevalent in small vent species previously thought to be non-symbiotic.

## Introduction

A number of marine invertebrates rely on nutritional symbioses with chemosynthetic bacteria, particularly – but not exclusively – those inhabiting deep-sea hydrothermal vents and hydrocarbon seeps. [1, 2]. In these environments, visually dominant assemblages of chemosymbiotic invertebrates, such as siboglinid tubeworms, bathymodioline mussels, alvinocaridid shrimps, or abyssochrysoidean snails flourish. They benefit the turbulent mix of cold, oxygenated seawater with reduced chemicals from vent and seep fluids (e.g., H_2_S, CH_4_, H_2_) [3] that creates ideal conditions for the energy metabolism of symbionts and the fixation of inorganic carbon [2]. A strong selection by host for one/a few bacterial lineages, often belonging to *Campylobacterota* and *Gammaproteobacteria*, is typically observed [4]. Because the chemical composition of fluids – in particular reduced compound concentrations – can strongly vary spatially and temporally, metabolic capacities of the symbionts constraint the ability of their hosts to inhabit distinct environments [5, 6]. Dual symbioses with different partners are often observed in some *Bathymodiolus* mussels or *Ifremeria* and *Gigantopelta* snails harbouring both sulfur-oxidizing and methanotrophic bacteria, enabling them to cope with the environmental variability of their habitats [7, 8]. Similarly, the metabolic flexibility of symbionts themselves can also enable the use of distinct energy sources, with thiotrophic symbionts being also hydrogenotrophic, as observed in some *Bathymodiolus* mussels or *Alviniconcha* snails [6, 9, 10].

Much of the research on vent holobionts has focused on iconic foundation species, with smaller species receiving considerably less attention. Only recently have chemosymbioses been discovered in a number of smaller, mixed diet macrofaunal species. Examples include *Sericosura* sea spiders from methane seeps of the eastern Pacific margin [11], *Ostiactis* anemones from hydrothermal vents in the Gulf of California [12] and *Rimicaris chacei* shrimps on the Mid-Atlantic Ridge [13] Furthermore, recent synthesis revealed a gradient of epibiotic bacterial colonisation across the family level among alvinocaridid shrimps, highlighting different trophic strategies and a gradual evolution towards greater reliance on symbiosis [14].

Peltospirid snails represent another rare vent-endemic lineage where numerous trophic strategies, including symbiotrophy, coexist within the same family. Most peltospirid snails are assumed to feed on microbial mats, but the use of endosymbiosis has been established in the large-bodied species *Gigantopelta aegis, G. chessoia*, and *Chrysomallon squamiferum* [15–17]. Unlike most chemosymbiotic molluscs, *Chrysomallon* and *Gigantopelta* species do not host their symbionts in their gills but within an enclosed part of their body, the oesophageal gland, which is greatly enlarged [16]. This is linked to other anatomical adaptations such as an enlarged circulatory system that supplies hydrogen sulfide and carbon dioxide to the endosymbiont [16, 18]. Despite superficial similarities, endosymbiosis in *Chrysomallon* and *Gigantopelta* is thought to have arisen convergently along their evolution [16]. While *Chrysomallon* hosts a single sulfur-oxidizing *Gammaproteobacteria* symbiont [19], *Gigantopelta* has two gammaproteobacterial partners, one methanotroph and one sulfur-oxidizing [17]. Post-settlement development is also distinct. A significant reorganization of internal anatomy (‘cryptometamorphosis’) and symbiont horizontal transmission occur in *Gigantopelta*. Conversely, an isometric growth with mixed symbionts transmission mode, both vertical and horizontal, characterize *Chrysomallon* [20, 21]. Symbiotic relationships have been suggested in *Hirtopelta* due to its exceptionally large gill containing numerous bacteriocytes, although unconfirmed yet via molecular methods. [22].

The discovery of two new vent fields on the Mid-Atlantic Ridge (MAR) in 2023, namely Hydra (25°N) and Falkor EMARK (23.5°N) [23], led to the findings of another potentially symbiotic peltospirid: *Peltospira gargantua* aggregating close to vent orifices [24]. Like the other chemosymbiotic peltospirids, this large species exhibits an enlarged oesophageal gland and circulatory system combined with a reduction of its alimentary canal [24]. The restricted geographic range of *P. gargantua* to these two vent fields containing high hydrogen gas concentrations, especially at Hydra (up to 60mM) [25], could be linked to specific metabolic requirement of their potential symbionts. Most other MAR vent fields between Moytirra (45.5°N) and Ashadze (13°N) are colonized by a smaller congener, *Peltospira smaragdina* also occurring close to vent emissions [26–28]. This species has been suggested to be pioneer settlers that graze on bacterial mats [26, 29, 30], yet the low δ^15^N is suggestive of putative symbiosis [31].

Here, we investigate if the two *Peltospira* snails inhabiting MAR vents host bacterial endosymbionts, integrating various approaches. Our questions are: 1) Do *P. gargantua* and/or *P. smaragdina* snails host chemosymbiotic bacteria, and if so, in which organs are they located? 2) Which bacterial lineages are hosted? 3) What are their metabolic potentials, and do they differ between the two hosts? 4) Do the two species share the same trophic strategies and degree of dependence on their endosymbionts?

## Material and methods

### Sample collection

We collected 32 *Peltospira* specimens from Lucky Strike, TAG, Hydra, Falkor EMARK, Snake Pit & Puy des Folles hydrothermal vent fields on the MAR (1700 m to 3993 m) (Figure 1 and Table S1). Specimens were sampled using suction samplers of the human-occupied vehicle (HOV) *Nautile* during the BICOSE 2 (https://doi.org/10.17600/18000004) and BICOSE 3 (https://doi.org/10.17600/18002399) expeditions on board R/V *Pourquoi pas?*, the remotely operated vehicle (ROV) Victor6000 during the Momarsat 2023 expedition (https://doi.org/10.17600/18002419) on board the R/V *L’Atalante* and the remotely operated vehicle (ROV) *SuBastian* during the expedition FKt230303 on board R/V *Falkor (too)* [23]. Immediately after collection, specimens were directly stored at −80°C, preserved in 80% ethanol for metagenomic and stable isotopes analyses or fixed on board for fluorescence in situ hybridization (FISH) analyses and transmission electron microscopy (TEM) (see below). Individual dedicated to anatomy study was fixed on board in 4% formalin.

**Figure 1.**
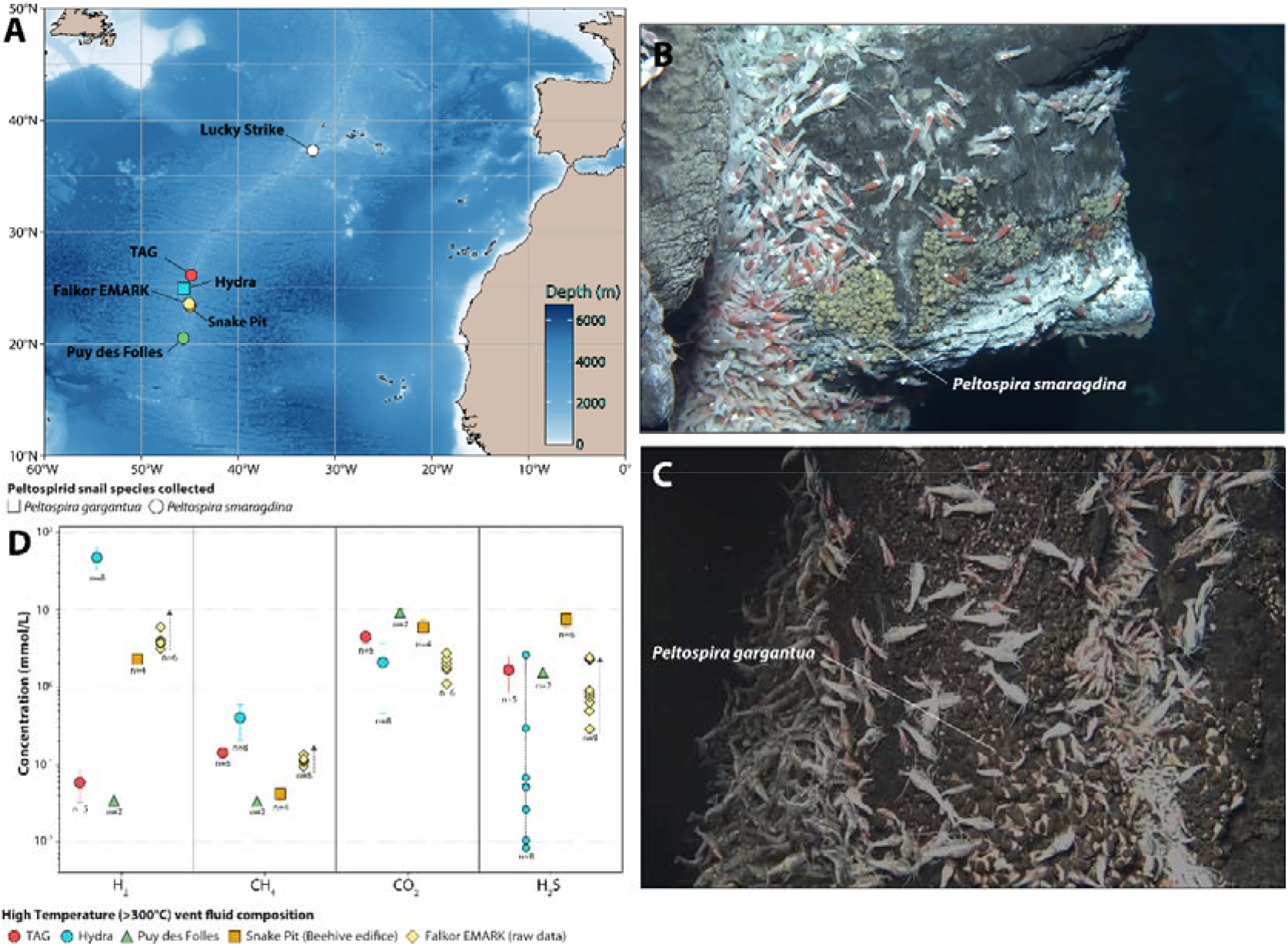
Geographic distribution and fluid geochemistry of hydrothermal vents inhabited by Peltospira holobionts **A.** Map of the hydrothermal vent fields from which *Peltospira gargantua* and *Peltospira smaragdina* were sampled along the Mid Atlantic Ridge (MAR) for this study. **B**. Patch of *Peltospira smaragdina* individuals surrounded by *Rimicaris* shrimps on the Beehive edifice at the Snake Pit vent field (credit: BICOSE 2 expedition). **C**. Colony of *Peltospira gargantua* individuals surrounded by *Rimicaris* shrimps at the Hydra vent field (credit: BICOSE 3 expedition). **D**. Dissolved gas concentrations (mmol/L) in high-temperature venting fluids from the different hydrothermal fields. For TAG, Snake Pit, Hydra and Puy des Folles, given concentrations represent endmember values (extrapolated to 0 Mg from n fluid samples). Error bar is 1 SD). At Hydra, H_2_S measurements are highly scattered, likely due to sulfide precipitation within the sampler and/or entrainment of chimney material during sampling. This prevents from reliable Mg correlations and endmember calculation. Individual measurements are therefore reported. For Falkor EMARK, major inorganic chemistry (including Mg) is not yet available, precluding endmember extrapolation; individual measurements from different samplings are shown instead.

We also collected high-temperature fluids using titanium “major” samplers for chemical characterization. Dissolved ΣH_2_S was determined on a separate aliquot prior to gas extraction by iodometric titration with triiodide, with residual triiodide measured on a spectrophotometer at 400 nm and ΣH_2_S derived from the equivalence volume. Gases (CO_2_, H_2_, CH_4_) were subsequently extracted on board using a vacuum line and quantified by FID- and TCD-GC following published methods [32].

### Three-dimensional reconstruction of internal anatomy

A specimen of *P. smaragdina* selected for micro-computed tomography (μCT) was contrast-stained in ethanol solution containing 0.3% phosphotungstic acid (PTA) and 3% dimethyl sulfoxide (DMSO) [33], then mounted while submerged in ethanol and scanned at the ANATOMIX beamline, Synchrotron SOLEIL (France). A polychromatic (“pink”) X-ray beam with a mean energy of 40 keV (filters: 26 μm Au, 100 μm Cu, 250 μm Dy) was combined with a Hasselblad optical system at 2.1× magnification. Images were acquired with a Hamamatsu ORCA Flash v4 detector at effective voxel size of 3.09 μm. Higher magnification scans (5×) with a Mitutoyo objective yielded a voxel size of 1.3 μm. Specimens were rotated over 180°, and 2,000 projections collected. Tomographic reconstruction was performed using PyHST2 [34]. The reconstructed volumes were imported into Dragonfly v.2022.2.0.1409 (Object Research Systems) for pre-processing (brightness and contrast adjustments) and segmentation followed by three-dimensional reconstruction and rendering. For *P. gargantua*, a μCT-based 3D reconstruction has already been published [24] and was used for comparisons with the new *P. smaragdina* model. Both models were post-processed using the Smooth Surface function in Amira v2019.2 to produce the final display models. Absolute and relative organ volumes were calculated using Amira v2019.2 for *P. smaragdina*, with corresponding data for *P. gargantua* taken from [24].

### Stable isotopes

We isolated pieces of oesophageal gland, gills, and foot tissues from frozen stored specimens – 3 *P. gargantua* from Hydra and 5 *P. smaragdina* from TAG – that we dried at 50°C in oven for 2 days and reduced in homogenous powder with a manual potter. We analyzed stable isotope using a continuous flow-elemental analysis isotope ratio mass spectrometry at the University of Liège (Belgium), using a vario MICRO cube C-N-S elemental analyser (Elementar Analysensysteme GMBH, Hanau, Germany) coupled to an IsoPrime100 isotope ratio mass spectrometer (Isoprime, Cheadle, United Kingdom). We used the widespread δ notation expressed in %0 for isotopic ratios [35] and the Δ notation for trophic shifts as in [14]. Analyses were based relative to international references and internal laboratory standards (see supplementary for additional details).

### Transmission electron microscopy

We fixed whole animals, shells removed, in a glutaraldehyde 2.5%-seawater solution kept at 4°C in seawater/sodium azide (NaN_3_) rinsing solution until use in laboratory. We dissected samples into body parts (gills, oesophageal gland, digestive gland, gonad) and postfixed in osmium tetroxide 1 %. We then dehydrated these in increasing ethanol series (50, 70, 95 and 100 %) before embedding in Epon resin (48 h, 60 °C). We cut sections using a Reichert–Jung ultramicrotome in semi-thin (600 nm) and t Thin (60 nm) sections. We stained the first with toluidine blue for observations with an Olympus BX 61 and mounted the second on copper grids, contrasted using uranyl acetate for observations with a HITACHI H7700 TEM, operated at 80 kV.

### Fluorescent in situ hybridization (FISH)

We fixed specimens for 2h in 3% paraformaldehyde, rinsed and stored in 1:1 PBS/Ethanol at −20°C for further analyses following existing protocols [36]. In brief, we embedded whole individuals without their shells in PEG resin (Sigma-Aldrich, Merck KGaA, Darmstadt, Germany), trimmed the resin bloc into 8-10 μm sections with a RM 2255 microtome (Leica Biosystems, Nussloch, Germany) and mounted sections on glass slides (Menzel-Gläser Superfrost Plus, USA). After resin removal with ethanol, we hybridized tissues at 46°C for 3h in a reaction mix containing the general eubacterial probe Eub338 [37] the negative control probe Non-Eub338 or the *Campylobacterota* specific probe Epsy914 [38] at concentrations of 0.5 μM. After the washing steps, we added a SlowFade Gold antifade reagent containing DAPI (Invitrogen). We observed tissues using a Zeiss Axio Imager.Z2 microscope equipped with the Apotome.2 and Colibri.7 light source, and capture images with the Zen software (Zeiss, Oberkochen, Germany).

### DNA extraction and sequencing

We extracted DNA from the oesophageal gland of three *Peltospira* individuals per locality including *P. smaragdina* from TAG, Falkor EMARK and Snake Pit and *P. gargantua* from Hydra using the DNeasy Blood & Tissue kit (Qiagen, Hilden, Germany) following the manufacturer’s instructions. Preparation of Illumina libraries and sequencing were done at the Integragen sequencing platform. We also extracted high molecular weight DNA from oesophageal gland of two additional specimens – 1 *P. gargantua* from Hydra and 1 *P. smaragdina* from TAG – using the SDS method [39]. Preparation of long-read libraries and sequencing were done on the PacBio Sequel II platform at Novogene (Tianjin, China).

### Metagenomic analysis

We trimmed adapter sequences from raw reads using bbduk from bbmap v38.57 (https://sourceforge.net/projects/bbmap/) for Illumina reads and HiFiAdapterFilt v3.0.0 [40] for PacBio HiFi reads. We also controlled quality and filter short-reads using illumina-utils v2.12 [41]. We assembled Illumina reads into contigs using Megahit v1.2.9 [42] with the metasensitive mode and a minimum contig length of 1000 bp and the PacBio HiFi reads using Hifiasm-meta v0.3.4 [43] with default settings. We clustered contigs from Illumina reads using BASALT v1.0 with the ‘‘more-sensitive’’ parameter [44] and manually refined the obtained Metagenome-Assembled Genomes (MAGs) with the “anvi-refine” program of Anvi’o v8_dev [45]. For PacBio reads, we clustered contigs with MaxBin v2.2.7 [46] and circularized symbiont genomes using Hifiasm-meta v0.3.4 or Flye v2.9. 3 [47] (see supplementary methods for additional details on workflows).

Combining the two MAGs datasets, we selected unique representative MAGs with ‘‘anvi-dereplicate-genomes’’ using the pyANI program, a similarity threshold of 98% average nucleotide identity (ANI), a coverage threshold of 30% and the Qscore method. Finally, we realized a final mapping of all metagenomes on the dereplicated MAGs to calculate their mean coverage and detection in each sample. We also assessed the completeness, contamination (redundancy) and genome size of these MAGs using CheckM2 v1.0.1 [48].

We obtained taxonomic affiliation of our MAGs using the classify workflow of GTDB-Tk v2.4.0 [49] and made multiple sequence alignments with taxa closely related to our MAGs using the GTDB-Tk “align” command. We trimmed multiple sequence alignments using TrimAl v1.4.1 [50] by removing positions with gaps in more than 50% of sequences. We reconstructed a maximum likelihood phylogenetic tree using IQ-TREE2 v2.2.5 [51] with the “WAG” general matrix model and 1000 ultrafast bootstrap replicates that we visualized using TreeViewer v.2.2.0 [52]. We calculated the pairwise ANI between our MAGs and their four most closely related bacterial references genomes from GTDB using “anvi-compute-genome-similarity” from anvi’o with the pyANI program. We used METABOLIC v4.0 [53] to estimate gene content and the completeness of various metabolic pathways.

### Phylogeny, synteny and protein structure prediction of hydrogenases

Annotated Ni-Fe hydrogenase sequences were classified into functional sub-groups using the HydDB database [54] and compared with reference sequences (see supplementary methods). We aligned sequences using MUSCLE v5.3 [55], and removed poorly aligned regions using TrimAl v1.4.1 [50]. A maximum-likelihood phylogenetic tree was reconstructed using IQ-TREE2 v3.1.0 [51], with the best-fit substitution model automatically selected by ModelFinder [56] and evaluated with 1,000 ultrafast bootstrap replicates. The resulting phylogenetic tree was visualized using Tree Visualization By One Table (tvBOT) [57].

Whole-genome synteny between the PacBio-assembled symbiont genomes was assessed using pairwise dot plot analysis implemented in JCVI v1.6.5 [58]. To investigate the conservation of Ni-Fe hydrogenase-associated regions, we performed microsynteny analysis between Ni-Fe genes which clustered within the same phylogenetic lineage using the MCScanX algorithm [59] implemented in JCVI and visualized using jcvi.graphics.synteny. In addition, the gene orders surrounding the two Ni-Fe hydrogenases in *P. gargantua* symbiont and the corresponding hydrogenase locus in *P. smaragdina* were compared to characterize differences in Ni-Fe hydrogenase gene organization. We predicted structures of Ni-Fe hydrogenases using AlphaFold3 [60] to evaluate structural similarity and conservation against known references (see supplementary methods).

## Results

### Fluid Geochemistry

Both Hydra and Falkor EMARK are located off-axis suggesting potential influence from ultramafic, mantle-derived lithologies [25]. At Hydra, endmember fluids (i.e. extrapolated to 0-Mg) contain exceptionally high H_2_ (~45 mM, and possibly up to 60 mM; [25]), but comparatively low H_2_S (<0.5 mM), CH_4_ (~0.4 mM), and ΣCO_2_ (~2 mM) (Figure 1D), the latter comparable to ambient seawater, suggesting limited magmatic input. EMARK similarly exhibits elevated H_2_ (≥4.6 mM), with CH_4_ ~0.1 mM, ΣCO_2_ ~2.6 mM, and H_2_S ~1–2 mM (Figure 1D). These concentrations were measured in a 360°C fluid without correction for seawater entrainment and therefore represent lower bounds for the endmember. In contrast, TAG and Snake Pit are located within the axial valley and hosted by neovolcanic basalt [25 and references therein]. TAG fluids are typical of basalt-hosted systems, with low H_2_ and CH_4_ (0.05 and 0.14 mM) and moderate ΣCO_2_ and H_2_S (4.5 and 1.7 mM). Despite its basaltic setting, Snake Pit exhibits elevated H_2_ (up to 3.4 mM), while CH_4_ remains low (~0.05 mM); ΣCO_2_ and H_2_S reach 6 and 7.6 mM, respectively, making Snake Pit the most CO_2_ and sulfur-rich of the four sites (Figure 1D).

### Comparative 3D anatomy

Both *Peltospira* species exhibited hypertrophied oesophageal glands, becoming pouch-like and extending posteriorly towards the visceral mass (Figure 2A-D). However, the oesophageal gland of *P. gargantua* (13.87% of body volume) is thicker dorsoventrally and is much more voluminous than *P. smaragdina* (5.43% of body volume) (Figure S1). Other aspects of the internal anatomy are generally similar in organisation but with several key differences (Figure 2A-D, Table S2 and Figure S2). The radula of *P. smaragdina* is an order of magnitude larger than that of *P. gargantua* (0.25% of body volume in *P. smaragdina*, 0.02% in *P. gargantua*). *P. smaragdina* intestine loops above the oesophagus, as opposed to below the oesophagus in *P. gargantua. P. smaragdina* gill, heart and ventricle (8.34%, 2.67% and 0.55% of body volume, respectively) are less voluminous than those of *P. gargantua* (15.83%, 4.49% and 2.21% of body volume, respectively).

**Figure 2.**
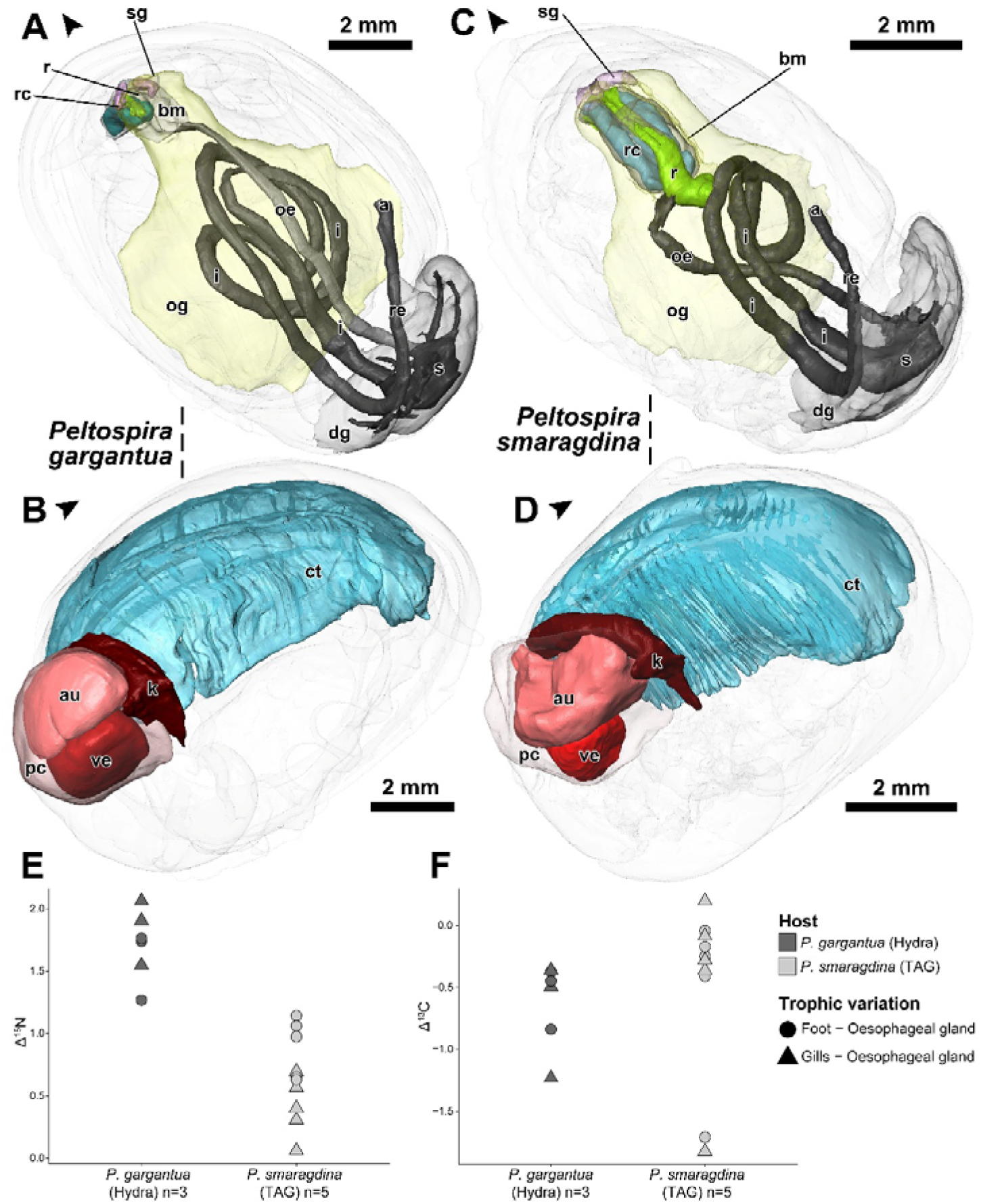
**A-B.** Three-dimensional anatomical reconstructions of *Peltospira gargantua* (reproduced from Chen et al. 2025) showing the digestive system **(A)** and the heart, kidney, and ctenidium **(B). C-D** Three-dimensional anatomical reconstructions of *Peltospira smaragdina* showing the digestive system (C) and the heart, kidney, and ctenidium (D). The body outline, buccal mass, ctenidium, oesophageal gland, pericardium, and digestive gland are shown in semi-transparency; arrowheads indicate the anterior of the animal. **E–F**. Trophic shifts between foot and oesophageal gland tissues (circle) or between gills and oesophageal gland tissues (triangle) for each individual of *P. gargantua* (dark grey) and *P. smaragdina* (light grey). **E**. Trophic shift for carbon isotopic ratios (Δ^13^C) **F**. Trophic shift for carbon isotopic ratios (Δ^15^N). Abbreviations: a, anus; au, auricle; bm, buccal mass; ct, ctenidium; dg, digestive gland; i, intestine; k, kidney (nephridium); oe, oesophagus; og, oesphageal gland; ov, ovary; pc, pericardium; r, radula; rc, radula cartilage; re, rectum; s, stomach; sg, salivary gland; ve, ventricle.

### Trophic analysis of Peltospira snails by stable isotopic composition

Isotopic ratios of analysed organs were comprised between −11.1 and −8.2 for δ^13^C, between 2.9 and 5.2 for δ^15^N and between 5.7 and 11.5 for δ^34^S (Table S3). We observed a δ^15^N and δ^34^S enrichments between the symbiont-hosting organ, the oesophageal gland, and the non-bearing symbiont organs (gills and foot) for both species (Figure 2E and S3) but no clear δ^13^C enrichment (Figure 2F). The trophic shift in nitrogen between organs was significantly higher for *P. gargantua* (Δ^15^N = 1.7 ± 0.3) compared to *P. smaragdina* (Δ^15^N = 0.6 ± 0.3; Wilcoxon tests, p > 0.05; Figure 6C) but no trophic shift was detected for carbon (Δ^13^C) or sulfur (Δ^34^S) (Wilcoxon tests, p > 0.05; (Figure 6D and S3).

### Structure and distribution of symbionts observed by FISH and TEM

All oesophageal gland samples showed abundant bacteria with FISH and TEM observations (Figure 3 and 4). Hybridization with the Eub338 yielded a positive signal – but not with the non-Eub338 control – in the oesophageal gland surrounding the oesophagus and the digestive system of *P. gargantua* (Figure 3A and 3C) and *P. smaragdina* (Figure 4A). These bacteria are grouped together in bacteriocytes, which are more or less densely occupied (Figure 3B and 4B). In some bacteriocytes, the bacteria are clustered together (Figure 3B), while in others they are loosely dispersed (Figure 3B). These bacteria are bacilli measuring approximately 1×2 µm (Figure 4C). Hybridization with the - specific probe Epsy914 indicate they belong to *Campylobacterota* both for *P. gargantua* (Figure 3D) and *P. smaragdina* (Figure 4D).

**Figure 3.**
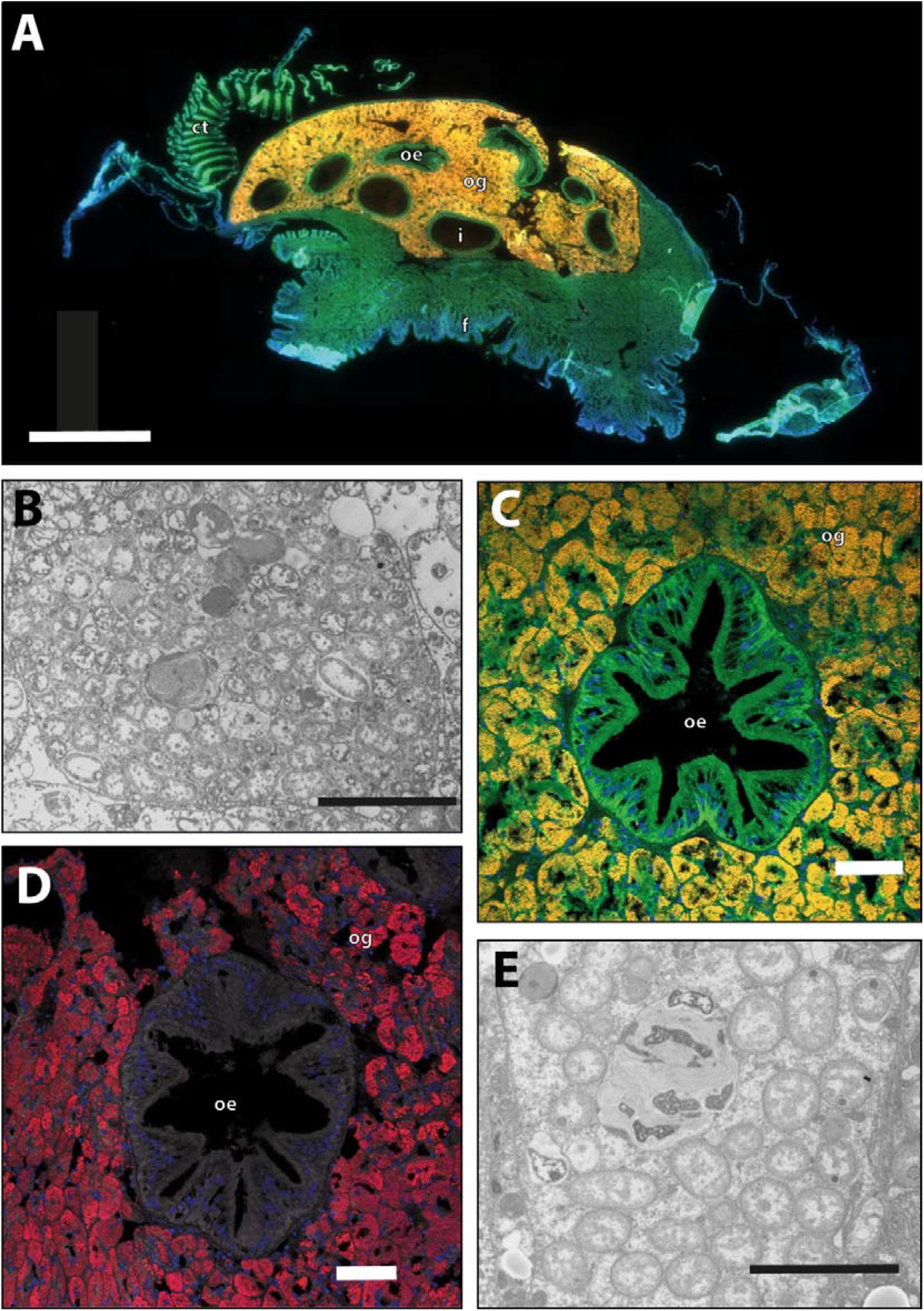
Fluorescence in situ hybridisation (FISH) **(A, C, D)** and Transmission electron microscopy (TEM) **(B, E)** observations of *Peltospira gargantua* snails showing bacterial endosymbionts in their oesophageal glands. oe: oesophagus; og: oesophageal gland; i: intestine; ct: ctenidium (gill); f: foot. **A.** Sagittal view of a whole *P. gargantua* individual hybridized with Eub338-Cy3 probe. Scale bar: 1 mm. **B**. *P. gargantua* bacteriocyte cells filled with bacilli-shaped endosymbionts in the oesophageal gland. Scale bar: 5 µm. **C**. Close-up view of *P. gargantua* oesophagus surrounded by the oesophageal gland filled with bacterial symbiont (Eub338-Cy3 probe). Scale bar: 50 µm. **D**. Close-up view of *P. gargantua* oesophagus surrounded by the oesophageal gland filled with *Campylobacterota* bacterial symbiont (Epsy914-Cy5 probe). Scale bar: 50 µm. **E**. *P. gargantua* bacteriocyte cells with bacterial endosymbionts and lysozyme in the oesophageal gland. Scale bar: 5 µm.

**Figure 4.**
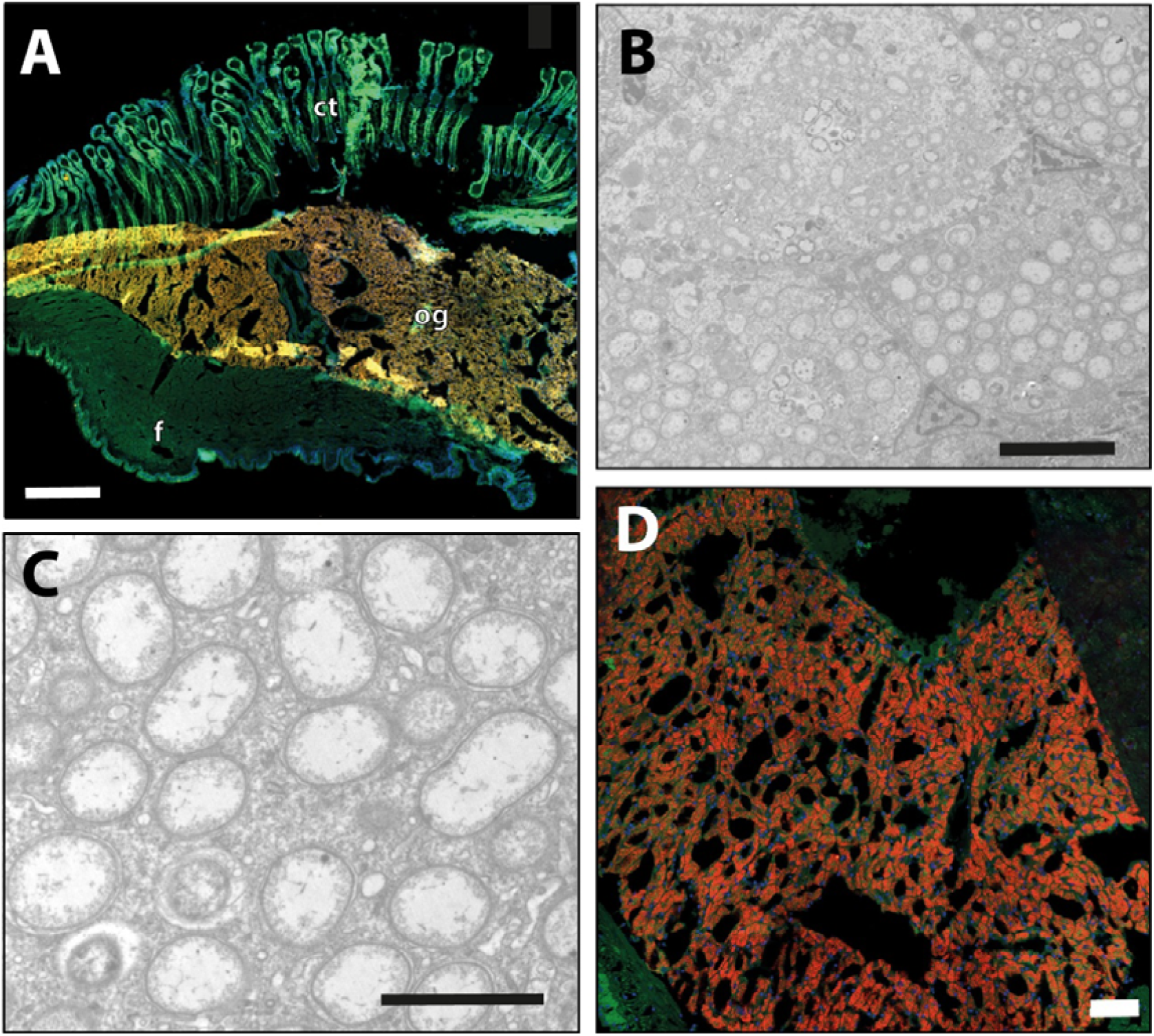
Fluorescence in situ hybridisation (FISH) **(A, D)** and Transmission electron microscopy (TEM) **(B, C)** observations of *Peltospira smaragdina* snails showing bacterial endosymbionts in their oesophageal glands. og: oesophageal gland; ct: ctenidium (gill); f: foot. **A.** Sagittal view of a *P. smaragdina* individual hybridized with Eub338-Cy3 probe. Scale bar: 400 µm. **B**. *P. smaragdina* bacteriocyte cells filled with bacilli-shaped endosymbionts in the oesophageal gland. Scale bar: 5 µm. **C**. Close-up view of *P. smaragdina* bacterial endosymbiont cells in the oesophageal gland. Scale bar: 2 µm. **D**. Close-up view of *P. smaragdina* oesophageal gland filled with *Campylobacterota* bacterial symbiont (Epsy914-Cy5 probe). Scale bar: 50 µm.

Within the bacteriocytes, numerous bacterial cell division can also be observed as well as numerous lysosomes, some containing degrading bacteria, indicating a possible process of bacterial digestion (Figure 3E). The same global organization is observed in the oesophageal gland of both species, with slightly more lysosomes/bacterial degradation in *P. smaragdina*. Many bacteria showed dense inclusions in the cytoplasm although more numerous in *P. gargantua* (Figure 3B and 3E) than in *P. smaragdina* (Figure 4B and 4C). Some bacteria are also present, scattered in the food bolus, along with numerous mineral particles (Figure S4). TEM observations of other tissues including gills, digestive glands, and gonads revealed no significant specific colonization by bacteria in either species (Figure S4). However, large clusters of bacteria were occasionally observed on the surface of the gill filaments in both gill samples taken from *P. smaragdina* (Figure S4).

### Genome sequencing, taxonomic affiliation & phylogenetic position of Peltospira symbionts

After assembly from Illumina reads, automatic and manual binning and dereplication at a 98% similarity threshold, we obtained a final collection of 8 high-quality MAGs (Figure 5A and Table S7) (see supplementary results for additional details). Taxonomic affiliation inferred from bacterial single-copy core genes (Table S8) showed the presence of one *Sulfurimonas* MAG per vent field (4 MAGs), two *Sulfurovum* MAGs isolated at Hydra and TAG, as well as two *Metamycoplasmataceae* MAGs isolated at Falkor EMARK (Figure 5A). The four *Sulfurimonas* MAGs were by far the most abundant MAGs, with a mean coverage of 3251.96 to 8323.98 for the different *Peltospira* individuals from the same field. In comparison, the two *Sulfurovum* MAGs could only be detected in *Peltospira* individuals from which they were isolated – Pga07 and Psm09 – with a mean coverage of 7.75 and 38.8 respectively. The two *Metamycoplasmataceae* MAGs also had a low abundance, with a mean coverage between 11.66 and 76.77 for the *Peltospira* individuals from Falkor EMARK.

**Figure 5.**
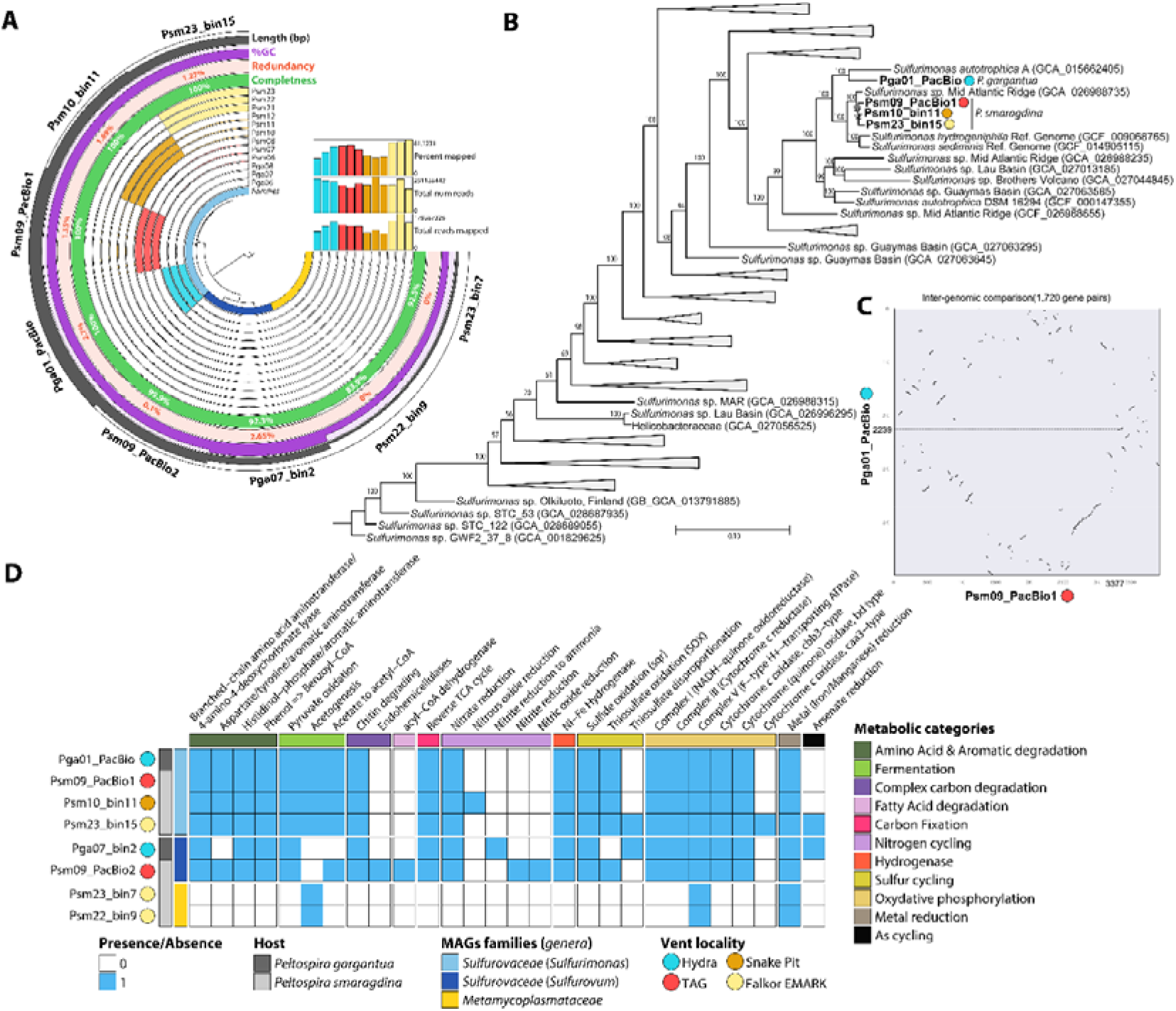
Metagenomic analyses of *Peltospira gargantua* and *P. smaragdina* oesophageal glands **A.** Static image from the anvi’o interactive display for the 12 *Peltospira* oesophageal gland samples with the 8 metagenome assembled genomes (MAGs) reconstructed in this study. From inner to outer layers: phylogenomic tree based on concatenated protein-coding genes according to GTDB-Tk, genome family and genus based on GTDB-Tk, 12 view layers with mean coverage information about MAGs across samples stored in the profile database – *: mean coverage of *Sulfurovum* and Metamycoplasmataceae MAGs are very low and cannot be distinguished –, percentage completeness, and redundancy according to CheckM2, GC-content and Genome lengths of each MAGs. The horizontal layers show the percentage of reads mapped, total number of reads mapped, and total number of reads for each sample **B**. Maximum likelihood phylogenetic tree using IQ-TREE v2.0.3 with the “WAG” general matrix model and 1000 bootstrap replicates visualized using TreeViewer for *Sulfurimonas* MAGs and their closest relatives. (see Figure S5 for *Sulfurovum* and Metamycoplasmataceae phylogenetic trees). **C**. Pairwise whole-genome dot plot analysis generated using JCVI, showing fragmented macro-syntenic relationships between *P. gargantua* symbiont from Hydra (Pga01_PacBio) and *P. smaragdina* from TAG (Psm09_PacBio1), with 1,720 homologous gene pairs distributed across discontinuous syntenic blocks. **D**. MAGs predicted functional capabilities according to METABOLIC showing presence/absence of key gene markers for different metabolic functions. For detailed results see Table S8 summarizing the output of METABOLIC analysis.

Our phylogenomic comparison revealed a clear divergence between the *Sulfurimonas* MAG of *P. gargantua* from Hydra and those of *P. smaragdina* from different vents, sharing 86.2 - 88.3 % ANI (Figure 5B and Table S9). Conversely, the *Sulfurimonas* MAGs of *P. smaragdina* from TAG, Snake Pit and Falkor EMARK were highly similar, sharing 97.0 - 97.5 % ANI between each other (Table S9). The two *Sulfurovum* MAGs, each found in a single individual, were clearly distinct from one another, sharing only 71.9% ANI whereas the two *Metamycoplasmataceae* MAGs from Falkor EMARK were relatively close from each other with 94.7% ANI (Figure S5 and Table S9) (see supplementary for details on closest known relatives).

Whole-genome pairwise macro-synteny analysis between the *Sulfurimonas* MAGs of *P. gargantua* and the one of *P. smaragdina* from TAG revealed highly fragmented syntenic relationships, with 1,720 homologous gene pairs distributed in scattered syntenic blocks rather than forming continuous collinear regions (Figure 5C), suggesting substantial genome rearrangement between the two symbiont genomes. Genome sequencing using PacBio HiFi reads have assembled two circular and representative symbiont genomes for both *Peltospira*, with the genome size of 3,406,356 bp for *P. gargantua* symbiont and 3,366,146 bp for *P. smaragdina* symbiont. The genome completeness and heterozygosity are 100% and 2.61% for *P. gargantua* symbiont and 99.99% and 1.56% for *P. smaragdina* symbiont. Given their high quality with circularized genomes, we propose two new candidate names for the main symbiont MAGs: *Candidatus* Sulfurimonas gargantua for the one hosted by *P. gargantua* and *Candidatus* Sulfurimonas smaragdina for the one hosted by *P. smaragdina*.

### Metabolic potential of symbionts

All MAGs affiliated to *Sulfurimonas* displayed very similar metabolic capabilities between the different vent fields with only minor differences of additional pathways in *Candidatus* Sulfurimonas smaragdina from Falkor EMARK and Snake Pit (Figure 5D). All *Sulfurimonas* MAGs encoded the entire set of genes to fix inorganic carbon trough the reverse TCA cycle – including the key acetate citrate lyase (acL), a complete electron transport chain gene set for oxidative phosphorylation, as well as a sulfide-quinone reductase gene (sqr) for sulfide oxidation and gene of the SOX pathway for thiosulfate oxidation (Figure 5D). Yet, *Candidatus* Sulfurimonas gargantua at Hydra differed from the different *Candidatus* Sulfurimonas smaragdina by having a truncated SOX pathway lacking SoxB and SoxC genes instead of the entire SoxABCDXYZ set (Table S10). The low-abundance MAGs affiliated to *Sulfurovum* exhibited metabolic capacities relatively similar to those of MAGs affiliated to *Sulfurimonas* whereas MAGs affiliated to *Metamycoplasmataceae* exhibited extremely limited metabolic capacities (Figure 5D).

All *Sulfurimonas* MAGs encoded Group I Ni-Fe hydrogenases for energy production based on HydDB assignment but differed in terms of copy numbers, with one Ni-Fe hydrogenase gene copy in the *Candidatus* Sulfurimonas smaragdina from TAG and Snake Pit, two gene copies in *Candidatus* Sulfurimonas gargantua and three gene copies in the *Candidatus* Sulfurimonas smaragdina from Falkor EMARK (Figure 6). Phylogenetic analysis of Group Ib Ni-Fe hydrogenases revealed that these sequences were divided into two major clades (Figure 6A). Notably, the different Ni-Fe hydrogenase copies encoded by *Candidatus* Sulfurimonas gargantua and *Candidatus* Sulfurimonas smaragdina from Falkor EMARK were positioned in different phylogenetic clades, indicating the presence of divergent Group 1b hydrogenase lineages within individual symbiont genomes.

**Figure 6.**
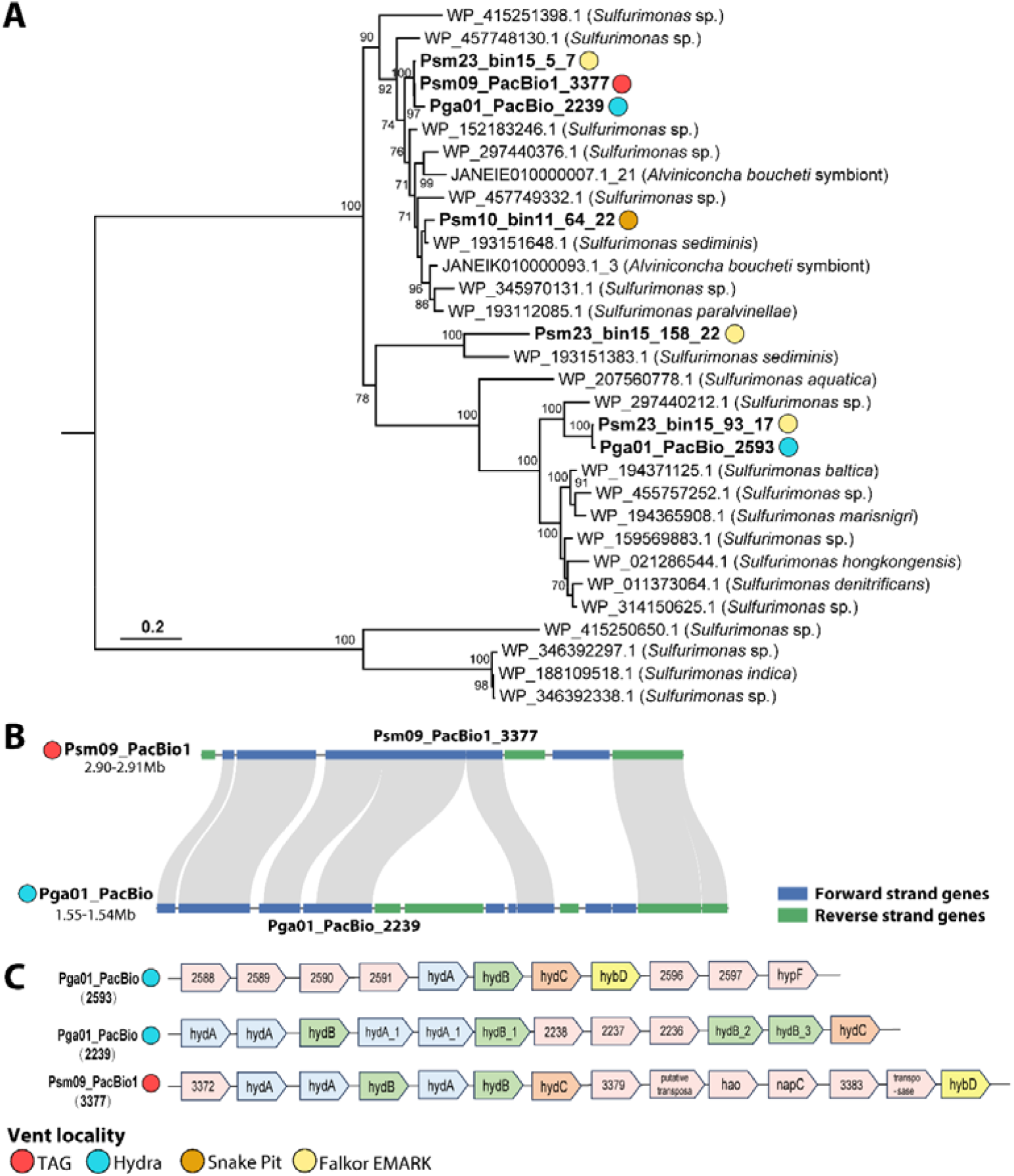
Phylogenetic and syntenic analyses of Ni-Fe hydrogenases. **A.** Maximum-likelihood phylogenetic tree of Group 1b Ni-Fe hydrogenases reconstructed using IQ-TREE2 with the ModelFinder Plus (MFP) strategy and 1,000 ultrafast bootstrap replicates. Group 2c Ni-Fe hydrogenases were included as the outgroup. Colored background strips indicate the ecological origins of the corresponding sequences. **B**. Microsynteny comparison of genomic regions surrounding the phylogenetically clustered hydrogenase loci Pga01_PacBio_2239 and Psm09_PacBio1_3377. Blue and green boxes indicate genes located on the forward and reverse strands, respectively. **C**. Comparative organization of Ni-Fe hydrogenase gene neighbourhoods. The genomic regions surrounding Pga01_PacBio_2239, Pga01_PacBio_2593, and Psm09_PacBio1_3377 were compared. The Pga01_PacBio_2239 locus contains truncated *hydA* and *hydB* genes and lacks a complete hydrogenase gene cluster, whereas the corresponding region in Psm09_PacBio1 contains intact *hydA, hydB*, and *hydC* genes with adjacent genes including transposase, *hao*, and *napC*. The Pga01_PacBio_2593 locus contains a complete hydrogenase gene cluster *(hydA*–*hydB*–*hydC*–*hybD*) together with the accessory gene *hypF*.

### Synteny analysis and structural comparison of symbiont hydrogenases

Microsynteny analysis showed that hydrogenase loci of *Candidatus* Sulfurimonas gargantua and *Candidatus* Sulfurimonas smaragdina from TAG were located within corresponding syntenic blocks (Figure 6B), but their gene neighbourhoods differed substantially.

The hydrogenase locus Pga01_PacBio_ 2239 region contained multiple truncated hydA and hydB genes but no complete hydrogenase gene cluster whereas the corresponding region in Psm09_PacBio1 contained complete hydA–hydB–hydC genes and additional transposase, hao, and napC genes (Figure 6C). The second locus in Pga01_PacBio (Pga01_PacBio_2593) exhibited a distinct genomic organization, with complete hydA–hydB–hydC–hybD cluster and the accessory gene hypF. (Figure 6C)

Structural comparison showed that the full-length hydrogenase encoded by most symbiont genomes shared a highly conserved overall fold with the Group Ib reference structure (1FRV). Across 511–526 aligned residues, these models exhibited TM-scores ranging from 0.96 to 0.98, RMSD values of 1.19–1.66 Å, and sequence identities of 42–49% within the aligned regions. In addition, all four conserved cysteine residues coordinating the Ni-Fe active site were correctly positioned in the predicted structures (Figure 7). Yet, Pga01_PacBio_2239 was C-terminally truncated and lacked the two conserved C-terminal cysteine residues required for Ni-Fe active-site coordination. Compared with the full-length reference structure, this model showed reduced structural similarity (TM-score = 0.66; 351 aligned residues). The loss of these conserved residues disrupts Ni-Fe active-site coordination and may affect the function of this hydrogenase.

**Figure 7.**
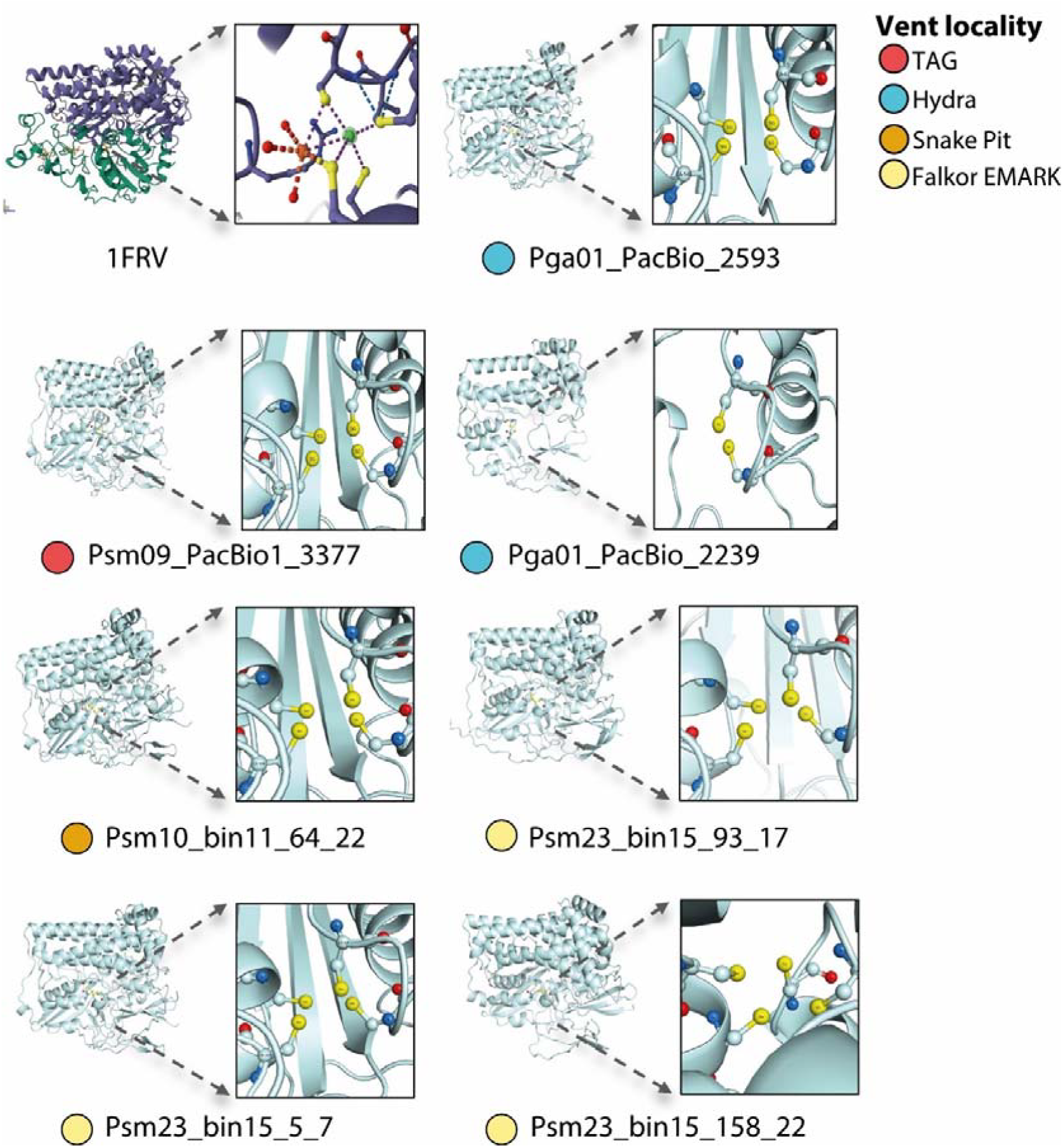
Structural prediction and comparison of Ni-Fe hydrogenases. AlphaFold3-predicted structures were aligned with the crystal structure of the Group 1b Ni-Fe hydrogenase from *Desulfovibrio gigas* (PDB: 1FRV). The Ni-Fe active center and conserved cysteine residues were visualized using PyMOL. Yellow spheres indicate the four conserved cysteine residues coordinating the Ni-Fe active site. Full-length hydrogenase models retained the conserved active-site architecture, whereas the truncated Pga01_PacBio_2239 model lacked the two conserved C-terminal cysteine residues due to loss of the C-terminal region.

## Discussion

*Peltospira smaragdina* has been known for over 30 years from hydrothermal vent sites along the northern MAR [28]. Yet, it has taken decades of accumulating indirect evidence [e.g., 31] and the recent discovery of its larger congener, *P. gargantua* [24], to hypothesise that *P. smaragdina* relies on bacterial endosymbionts for nutrition [31]. Our results supports this hypothesis and places both *P. gargantua* and *P. smaragdina* with the already known peltospirid holobionts, *Chrysomallon squamiferum* and the two *Gigantopelta* species, highlighting that chemosymbiosis might be more widespread in this family than previously thought [15–17]. Since *P. smaragdina* and *P. gargantua* were sister species based on the phylogenetic analysis [24], the ancestor of these two species was likely chemosymbiotic. This is another case of convergent evolution of chemosymbiosis in the Peltospiridae besides *Chrysomallon, Gigantopelta*, and potentially *Hirtopelta*.

### Peltospira snails host distinct Sulfurimonas symbiont lineages with potential local adaptation

Unlike other chemosymbiotic peltospirids with *Gammaproteobacteria* symbionts [17, 19], our results show that *P. smaragdina* and *P. gargantua* host *Campylobacterota* symbionts belonging to *Sulfurimonas* within bacteriocytes of their oesophageal gland, respectively named *Candidatus* Sulfurimonas smaragdina and *Candidatus* Sulfurimonas gargantua. This adds Peltospiridae to the list of chemosymbiotic animals, capable of hosting both *Gammaproteobacteria* and *Campylobacterota* symbionts, as previously observed in *Alviniconcha* snails and *Rimicaris* shrimps [8, 61].

Our results also show that these two *Peltospira* species host different *Sulfurimonas* lineages as their main symbiont (86.2 to 88.3% shared ANI). This suggests bacterial partner choice differs by host species similar to previous observations of species-specific symbionts in *Chrysomallon* and *Gigantopelta*, despite co-occurring side-by-side [17]. As seen in *P. smaragdina*, symbionts within one host species were very similar across different localities (97% shared ANI) suggesting that they belong to the same bacterial species, with different local strains at each vent. This mirrors the situation in *Chrysomallon squamiferum* [21], chemosymbiotic mussels and *Alviniconcha* snails [62–64]. The presence of other bacterial lineages in much lower abundance – *Sulfurovum* and *Metamycoplasmataceae* – imply that they represent facultative symbionts in the oesophageal gland or environmental bacteria as observed by TEM that pass through the oesophagus.

*Sulfurimonas* partners of *P. smaragdina* and *P. gargantua* exhibit very similar metabolic potentials, with an ability for both thiotrophy (sqr gene and the SOX system) and hydrogenotrophy (group 1 Ni-Fe hydrogenases). Flexibility of these holobionts to geochemical variations of the different sites they occupy along the MAR could thus be linked to variations in symbiont gene expression, favouring one energy pathway over another depending on the conditions, similar to observations and experiments on *Alviniconcha* snails and *Bathymodiolus* mussels [6, 9, 10]. Nevertheless, we found variations in the copy number of Ni-Fe hydrogenases among local symbiont populations of the two *Peltospira* species. This includes one gene copy in *Sulfurimonas* symbionts from TAG and Snake Pit, two for those from Hydra and three for those from Falkor EMARK. Moreover, the two hydrogenases copies of *P. gargantua* symbionts from Hydra also exhibited distinct genomic organization highlighting potential local adaptation to vent fluids with the highest H_2_ concentrations [25, 32]. Similarly, the presence of three hydrogenase copies in symbionts from Falkor EMARK despite similar H_2_ concentrations as Snake Pit, could be driven by the low concentrations of H_2_S at this vent field [25, 32]. Whether these hydrogenase copies are expressed and functional – particularly the one featuring a truncated active site – remain to be tested. Further transcriptomic analyses, combined with on board incubation experiments, comparing the two species across different geochemical contexts, will be required to better understand their physiological adaptations.

### Different levels of symbiosis dependence between sister Peltospira species

Anatomical features and stable isotopic compositions provide insights into the feeding ecology of both species and strongly suggest that *P. smaragdina* is less reliant on its chemosymbionts than *P. gargantua*. The symbiont-hosting oesophageal gland is nearly three times smaller in *P. smaragdina* than *P. gargantua*, as are the ventricle and the gill, which are half the size. An enlarged oesophageal gland with a higher degree of reliance on symbiosis is a trait seen across chemosymbiotic animals in order to increase the habitable space of the symbionts and therefore raising the energetic yield [2, 65, 66]. Similarly, a larger, muscular ventricle is considered to be an adaptive trait to meet the increased demand for oxygen by symbiotic bacteria which carries out oxidation reaction; a feature known from both other symbiotic peltospirids [18] and *Alviniconcha* and *Ifremeria* [65] snails. Increased gill volume despite a lack of gill symbionts is also seen in other chemosymbiotic peltospirids, especially in the scaly-foot snail, and is thought to be an adaptation to extract more oxygen which is often low in vent fluids [18]. The smaller body size of *P. smaragdina* also suggests a lesser reliance on chemosymbiosis as it often leads to gigantism [2]. Moreover, the radula is larger in *P. smaragdina* which is similar in size to non-symbiotic peltospirids [24], suggesting this species may still rely on the feeding apparatus. Conversely, *P. gargantua* radula is very reduced as observed in *Chrysomallon squamiferum* and *Gigantopelta* species [16, 18]. As such, *P. smaragdina* is likely a mixotrophic species that only relies partially on its symbionts, similar to the Atlantic alvinocaridid shrimp *Rimicaris chacei* [66, 67].

Trophic shift supports the anatomical features interpretation with a higher enrichment in nitrogen for *P. gargantua* suggesting a stronger reliance on its symbionts as nutritional source than *P. smaragdina*. The δ^13^C values of both species are clearly within the range of carbon sources fixed by rTCA, consistent with the metabolic potential of their symbionts [68] whereas their relatively low δ^15^N values also indicate limited ^15^N-enrichment to higher trophic levels [69], typical of symbiotroph species. Our results add to previous work on vent and seep mussels and shrimps [14, 70, 71], suggesting that mixotrophic diets, partially dependent on chemosymbiosis, are a more widespread feeding strategy than previously thought and that many such cases remain to be discovered.

## Acknowledgements

We thank the captain and crew of R/Vs *Pourquoi pas?* and *Falkor (too)* during the research cruises BICOSE2, BICOSE3, Momarsat 2023 and FKt230303, respectively, and extend this to the pilots and technical teams of HOV *Nautile* and ROVs Victor6000 and *SuBastian*. We also acknowledge chief scientists of these four cruises, Marie-Anne Cambon (Ifremer, France), Marjolaine Matabos (Ifremer, France) and David A. Butterfield (NOAA PMEL and University of Washington, USA) for their efforts in organizing and executing the expeditions. We also thank Dr. Olivier Rouxel (Ifremer) for the in-situ H_2_S measurements in hydrothermal vent fluids realized on board the R/V *Pourquoi pas?* TEM was performed at the electron microscopy platform of National Museum of Natural History (MNHN, Paris) with the help of Bérénice Piquet.

