## Supplementary Materials & Figures for "Two new gastropod holobionts from Atlantic hydrothermal vents showing different levels of endosymbiosis dependence"

### Supplementary materials & methods

#### Stable isotopes

For isotopic analyses, we used the following international references: Vienna Pee Dee Belemnite (for carbon), atmospheric air (for nitrogen) and Vienna Canyon Diablo Troilite (for sulfur). IAEA (International Atomic Energy Agency, Vienna, Austria) certified reference materials sucrose (IAEA-C-6; δ^13^C = −10.8 ± 0.5‰; mean ± s.d.), ammonium sulphate (IAEA-N-2; δ^15^N = 20.41 ± 0.12‰; mean ± s.d.) and barium sulphate (IAEA-SO5, δ^34^S = 0.5 ± 0.2‰) were used as primary analytical standards. Sulfanilic acid (Sigma-Aldrich; δ^13^C = −25.6 ± 0.4‰; δ^15^n = −0.13 ± 0.4‰; δ^34^S = 5.9 ± 0.5‰; means ± s.d.) was used as a secondary analytical standard; s.d. on multi-batch replicate measurements of secondary and internal lab standards (seabass muscle) interspersed with samples (one replicate of each standard every 15 analyses) were 0.2‰ for δ13C, 0.1‰ for δ^15^N and 0.3‰ for δ^34^S.

Trophic shift in carbon (Δ^13^C) was defined as the δ^13^C isotopic ratio of foot or gill tissues (the ‘’consumer’’) from which was subtracted the δ^13^C isotopic ratio of the oesophageal gland colonized by symbionts (the ‘‘nutrition source’’). Following the same principle, Δ^15^N and Δ^34^S were calculated as follows for each individual: Δ^15^N = δ^15^N_foot/gill_ − δ^15^N_gland_ and Δ^34^S = δ^34^S _foot/gill_ − δ^34^S_gland_. Statistical comparison of trophic shifts between the two *Peltospira* species were performed with Wilcoxon tests in a R v.4.4.3 statistical environment [1].

#### Library preparation for DNA sequencing

Illumina libraries were prepared at the Integragen sequencing platform with the Twist Library Preparation EF Kit 2.0 using 400 ng of DNA products. After enzymatic fragmentation, end-repair, A-tailing, ligation to UDI adapters full-length Illumina adapters, libraries were purified and sized with SPRI beads. Sized libraries were then quantified by qPCR. The sequencing was performed on an Illumina NovaSeq X Plus sequencer as paired end 2x150 bp reads.

For high molecular weight DNA extracted with the SDS methods, high-quality DNA (main band >30 kb) were used to construct HiFi SMRTbell libraries with the SMRTbell Express Template Prep Kit 3.0 (Pacific Biosciences). The constructed libraries were sequenced on the PacBio Sequel II platform at Novogene (Tianjin, China), with high-quality HiFi reads generated on the circular consensus sequencing (CCS) mode.

#### Metagenomic workflow

For manual refinement of MAGs obtained from BASALT [2], we generated using the metagenomic snakemake workflow implemented in Anvi’o v8_dev [3], an anvi’o contigs database from assembled contigs based on the identification of single-copy core genes with HMMER v3.3.2 [4] and open reading frames with Prodigal v2.6.3 [5], along with the annotation of gene functions with the KOfam HMM database of Kyoto Encyclopedia of Genes and Genomes (KEGG) orthologs [6, 7] and the NCBI Clusters of Orthologous Groups (COG) [8]. We also mapped short reads from each metagenome to the contigs using Bowtie2 v2.5.1 [9] and converted SAM to BAM files using samtools v1.18 [10]. In parallel, we profiled the BAM files using “anvi-profile” before combining all profiles into a single anvi’o profile with “anvi-merge.”

To obtain the circular-level genome of the symbiont, HiFi reads corresponding to the target bins were retrieved by mapping reads to the binned genomes using minimap2 v2.26 [11]. Then, these mapped HiFi reads were subsequently reassembled using Hifiasm-meta v0.3.4 or Flye v2.9. 3 [12] to generate circularized genomes.

#### Phylogeny, synteny and protein structure prediction of hydrogenases

Ni-Fe hydrogenase were initially annotated using METABOLIC v4.0 [13]. Functional annotations were further assigned by searching against the KEGG database using BLASTp v2.14.0 [14], the Clusters of Orthologous Groups (COG) database using eggNOG-mapper v4.5 [15], and the Pfam database using HMMER v3.4 [4]. For phylogenetic analyses, reference Group 1b Ni-Fe hydrogenase sequences from *Sulfurimonas* species were retrieved from HydDB, while Group 2c Ni-Fe hydrogenases were included as the outgroup. To provide additional reference sequences for phylogenetic analysis, genomes of *Alviniconcha boucheti* symbionts available from NCBI were annotated using the same workflow.

Based on predicted structures of Ni-Fe hydrogenases with AlphaFold3, structural similarity was evaluated by pairwise structural alignment against the crystal structure of the Group 1b Ni-Fe hydrogenase from Desulfovibrio gigas (PDB: 1FRV) [16] using the structural comparison tools available in the RCSB Protein Data Bank [17]. Structural conservation was assessed based on the root mean square deviation (RMSD) and TM-score values [18]. Conserved cysteine residues involved in Ni-Fe active-site coordination were identified based on sequence alignments and structural comparison with the reference hydrogenase structure. The spatial distribution of the four conserved cysteine ligands was visualized using PyMOL v3.1.8 [19], with the Ni-Fe active center and coordinating residues highlighted to examine the conservation of the catalytic architecture.

### Supplementary results

#### Metagenomic analyses of Sulfurimonas MAGs

Shotgun sequencing of total DNA recovered from the oesophageal gland of 12 *Peltospira* snails resulted in 1.55 billion reads with an average of 119 million reads per metagenome, of which 86.5% passed quality check on average (Table S4). Assembly of each sample resulted in 169K to 36K contigs longer than 1 kbp (Table S5). Our combination of automatic binning with BASALT and manual refined binning with Anvi’o resulted in 15 high-quality MAGs with low redundancy – below 3% – and high completeness – above 75% including 13 MAGs above 90%. HiFi sequencing of two additional *Peltospira* snails generated three circular-level genomes of the symbionts. Dereplication at a 98% similarity threshold eliminated 10 MAGs, grouping those from different individuals but originating from the same vent field together (Table S6).

Closest relative to the *Sulfurimonas* MAG of *P. gargantua* with 85.6% ANI was a *Sulfurimonas autotrophica* MAG isolated from the hydrothermal plume of the Von Damm vent field (Mid-Cayman Spreading Center) whereas closest relative to *Sulfurimonas* MAGs of *P. smaragdina* with 95.5 – 95.6 % ANI was a *Sulfurimonas* MAG isolated from hydrothermal fluid at the Rainbow vent field (MAR) (Figure 5B and Table S9). Closest relative to the two *Sulfurimonas* MAGs was a *Sulfurovum* isolated from the Longqi vent field (Southwest Indian Ridge) with 78.1 % ANI for Pga07_bin2 (*P. gargantua*; Hydra) and a *Sulfurovum* isolated from the Lucky Strike vent field (MAR) with 93.8 % ANI for Psm09_Pacbio2 (*P. smaragdina*; TAG). Closest relative of the two Metamycoplasmataceae MAGs with 72.2 - 72.4 % ANI was a Metamycoplasmataceae isolated from a microbial mat in the Guaymas basin (Gulf of California) (Figure S5 and Table S9).

**Supplementary Figures**

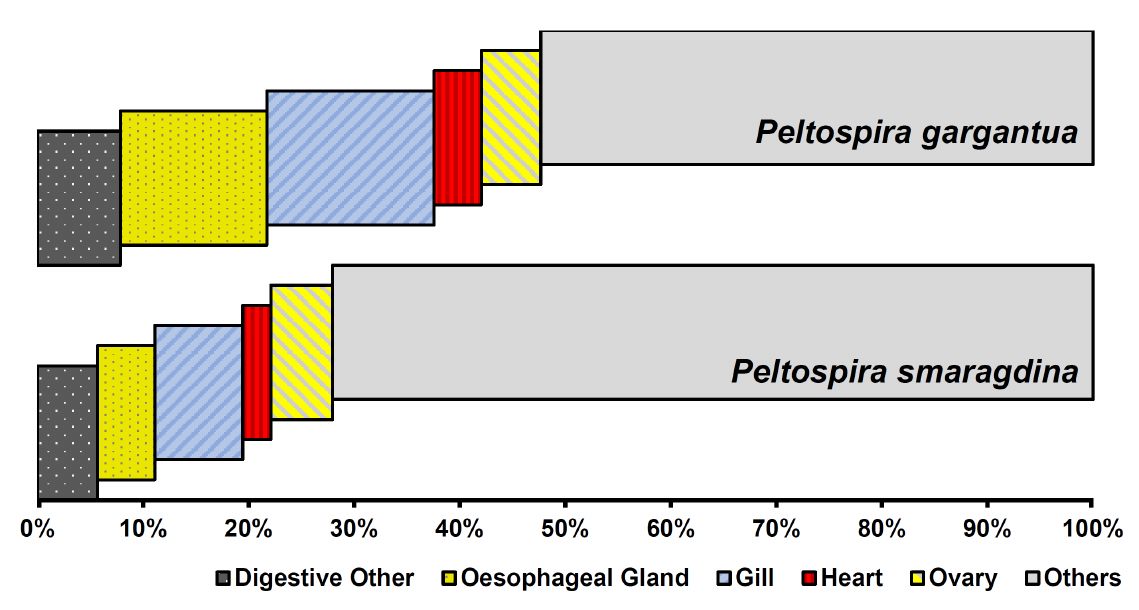

**Figure S1.** Relative percentage body volume of organs and organ systems in *Peltospira gargantua* and *P. smaragdina*, showing volume of the oesophageal gland, all other parts of the digestive system, gill, heart (auricle + ventricle), and the ovary, in relation to all other soft parts.

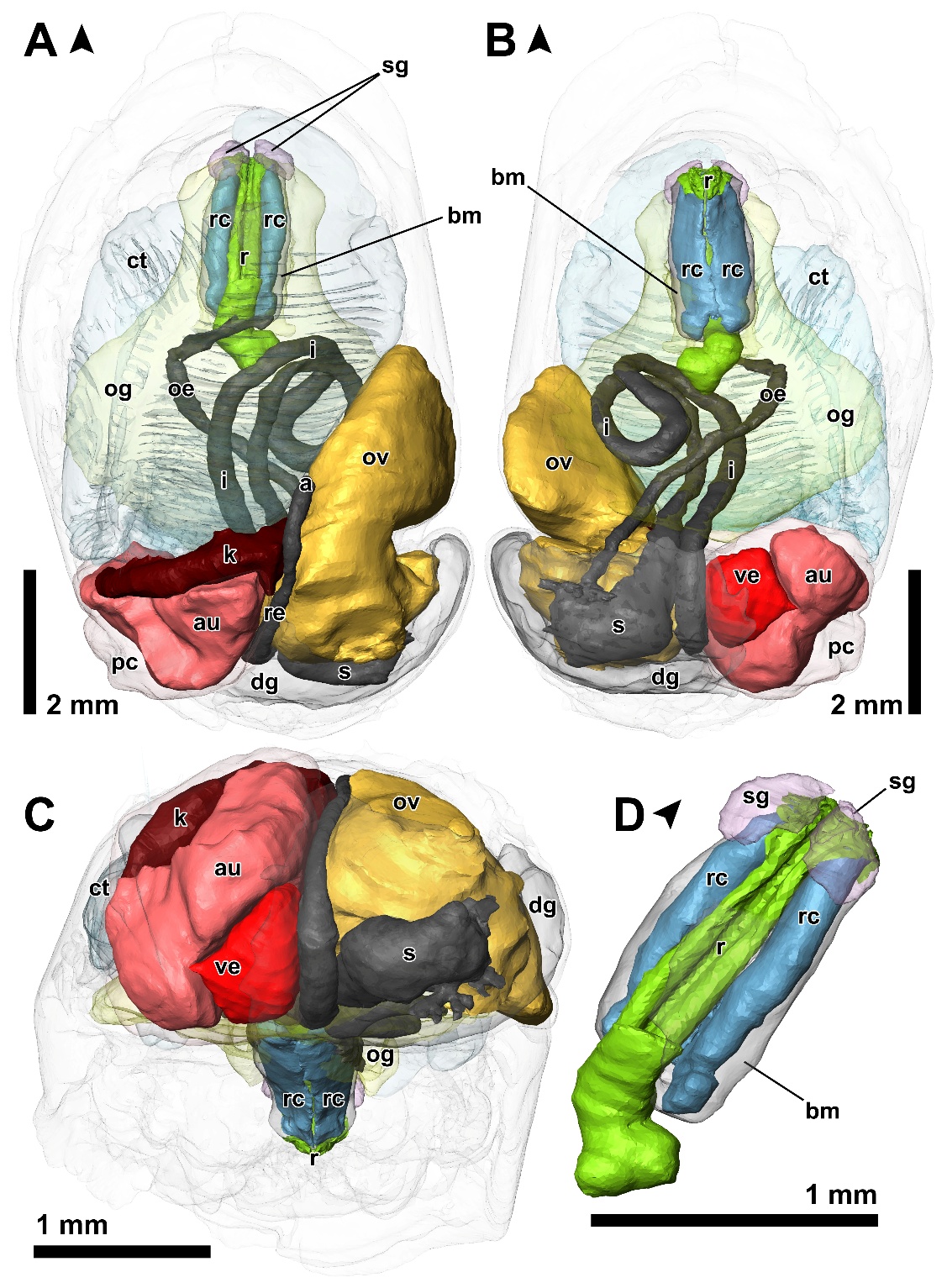

**Figure S2.** Three-dimensional anatomical reconstructions of *Peltospira smaragdina* showing the positions of major organs. The buccal mass, ctenidium, oesophageal gland, pericardium, and digestive gland are shown in semi-transparency. Arrowheads indicate the anterior of the animal. A, dorsal view. B, ventral view. C, rear view. D, enlarged view of the buccal mass showing the radula. Abbreviations: a, anus; au, auricle; bm, buccal mass; ct, ctenidium; dg, digestive gland; i, intestine; k, kidney (nephridium); oe, oesophagus; og, oesphageal gland; ov, ovary; pc, pericardium; r, radula; rc, radula cartilage; re, rectum; s, stomach; sg, salivary gland; ve, ventricle.

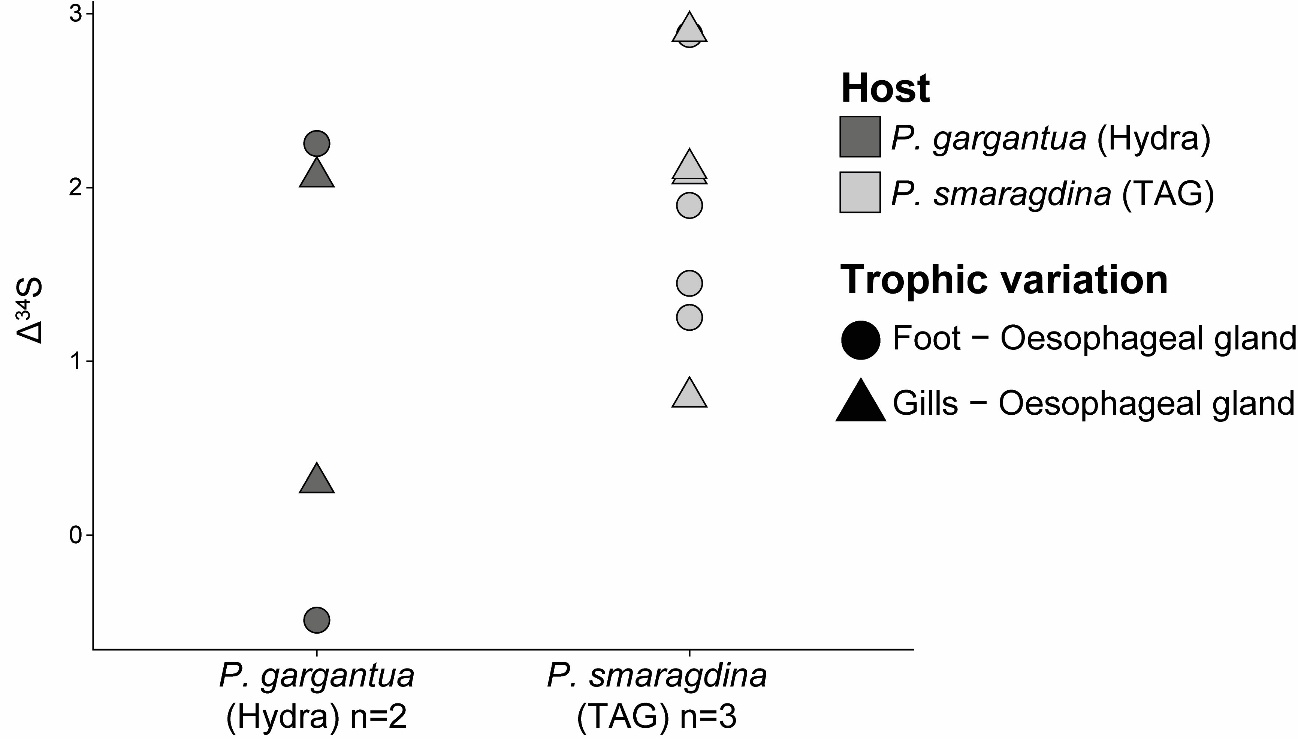

**Figure S3.** Trophic variations for sulfur isotopic ratios (Δ^34^S) between foot and oesophageal gland tissues (circle) or between gills and oesophageal gland tissues (triangle) for each individual of *P. gargantua* (dark grey) and *P. smaragdina* (light grey).

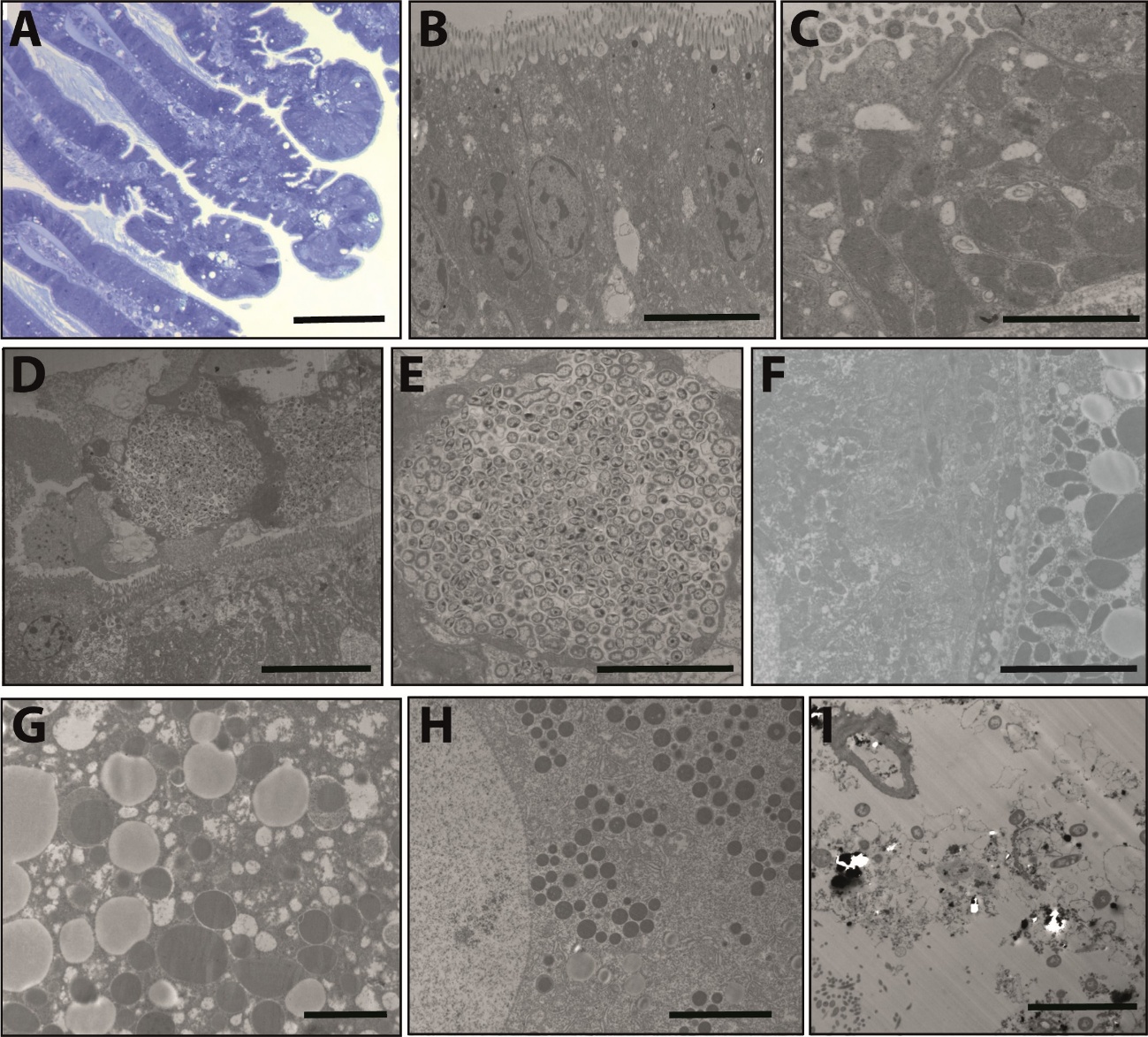

**Figure S4.** Additional light microscopy (**A**) and transmission electron microscopy (B-I) observations of *Peltospira* snails A. *P. gargantua* gills stained with toluidine blue. Scale bar: 50 µm. B. *P. gargantua* gill cells showing an absence of endosymbiotic bacteria. Scale bar: 5 µm. C. *P. smaragdina* gill cells showing an absence of endosymbiotic bacteria. Scale bar: 2 µm. D. Cluster of epibiotic bacteria on the surface of *P. smaragdina* gills. Scale bar: 10 µm. E. Close-up view of bacteria clusters at the surface of *P. smaragdina* gills. Scale bar: 5 µm. F. *P. gargantua* cells without endosymbiotic bacteria in the digestive gland (left side) or the gonads (right side) Scale bar: 5 µm. G. Absence of endosymbiotic bacteria in the gonad cells of *P. smaragdina*. Scale bar: 2 µm. H. Absence of endosymbiotic bacteria in the digestive gland cells of *P. smaragdina*. Scale bar: 5 µm. I. Bacterial cells in the alimentary bolus of *P. smaragdina*. Scale bar: 5 µm.

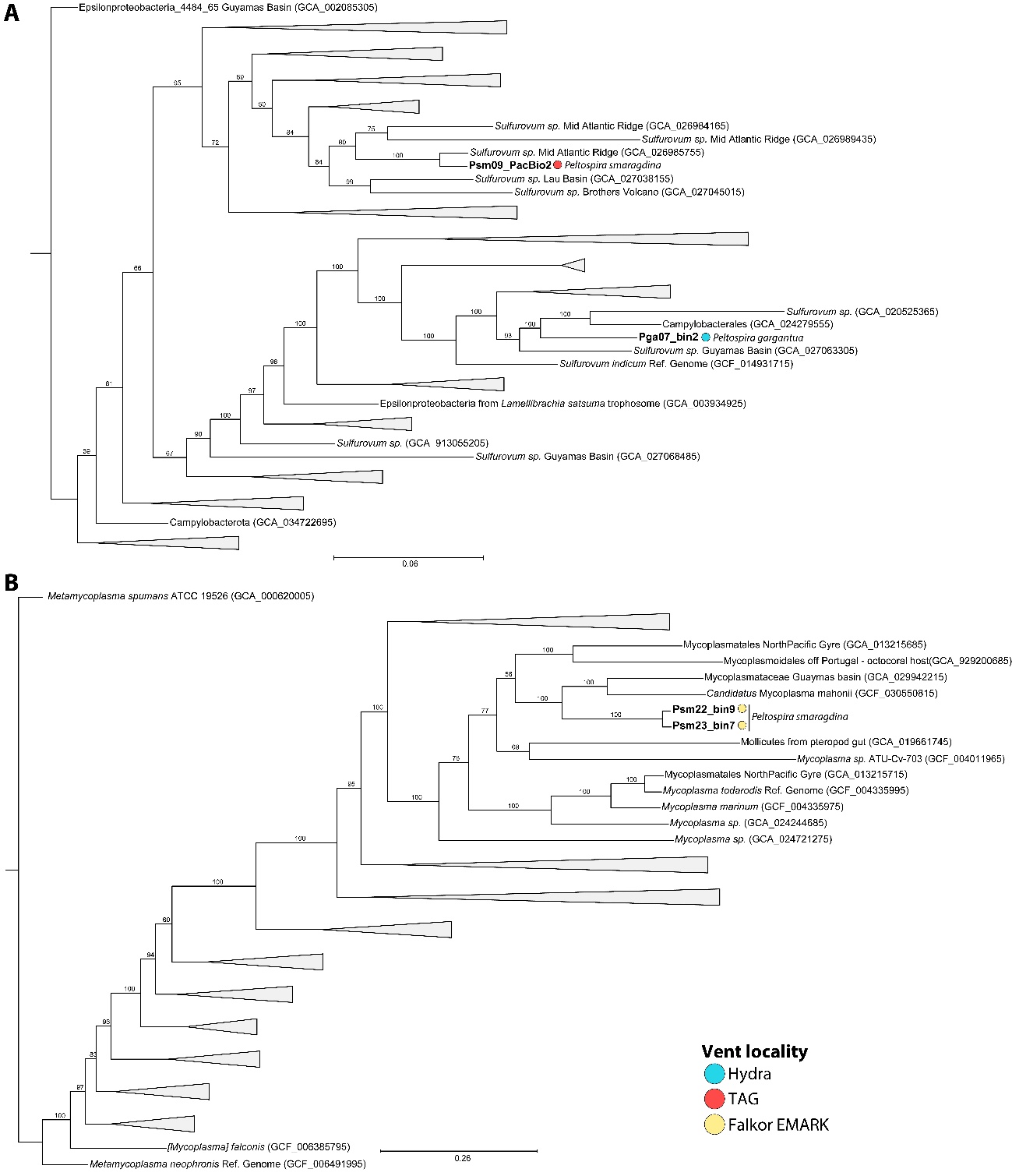

**Figure S5.** Maximum likelihood phylogenetic tree using IQ-TREE v2.0.3 with the “WAG” general matrix model and 1000 bootstrap replicates visualized using TreeViewer for *Sulfurovum* and *Metamycoplasmataceae* MAGs and their closest relatives.
